# Cutaneous Wounds in Mice Lacking Tumor Necrosis Factor-stimulated Gene-6 Exhibit Delayed Closure and an Abnormal Inflammatory Response

**DOI:** 10.1101/676411

**Authors:** Sajina Shakya, Judith A. Mack, Minou Alipour, Edward V. Maytin

**Author notes:** Corresponding author: Edward V. Maytin, MD, PhD, Department of Biomedical Engineering, Cleveland Clinic, Mailstop ND-20, 9500 Euclid Avenue, Cleveland, OH 44195, USA.

## Abstract

We investigated how loss of tumor necrosis factor-stimulated gene-6 (TSG-6) affects wound closure and skin inflammation. TSG-6 has several known biological functions including enzymatic transfer of heavy chain proteins (HC) from inter-α-trypsin inhibitor to hyaluronan (HA) to form HC-HA complexes. TSG-6 and HC-HA are constitutively expressed in normal skin and increase post-wounding, but are completely absent in TSG-6 null mice. Wound closure rates are significantly delayed in TSG-6 null mice relative to wildtype mice. Neutrophil recruitment is delayed in early wounds (12 h and Day 1), whereas late wounds (Day 7) show elevated neutrophil accumulation. <u>In addition, the granulation phase is delayed, with persistent blood vessels and reduced dermal collagen at 10 days. The pro-inflammatory cytokine TNFα is elevated >3 fold in unwounded TSG-6 null skin, and increases further after wounding (from 12 h to 7 days) before returning to baseline by day 10. Other cytokines examined such as IL-6, IL-10, and MCP-1 showed no consistent differences</u>. Importantly, reintroduction of TSG-6 into TSG-6 null wounds rescues both the delay in wound closure and the aberrant neutrophil phenotype. In summary, our study indicates that TSG-6 plays an important role in regulating wound closure and inflammation during cutaneous wound repair.

## INTRODUCTION

Cutaneous wound healing is a complex process that involves inflammation, keratinocyte/fibroblast proliferation, migration, angiogenesis, extracellular matrix (ECM) deposition and remodeling, all orchestrated by the specific actions of cytokines, chemokines, growth factors, and enzymes (Barrientos et al., 2008, Clark, 2001, Martin, 1997, Singer and Clark, 1999). One of these enzymes is Tumor necrosis factor (TNF)-stimulated gene-6 (*TSG-6*) protein, also known as TNF-α induced protein 6 (Tnfip6 or Tnfaip6). *TSG-6*, first identified by Lee et al. (Lee et al., 1990) in human foreskin fibroblasts stimulated with the pro-inflammatory cytokine TNFα, encodes a secretory glycoprotein with a molecular weight of ∼30-35 kDa (Milner and Day, 2003, Mittal et al., 2016, Wisniewski et al., 1993).

Functionally, TSG-6 appears to play important roles in ECM modification and in tissue inflammation. One of its important enzymatic activities is to catalyze the covalent transfer of heavy-chain (HC) protein from inter-alpha-trypsin inhibitor (IαI) and pre-α-inhibitor (PαI) to hyaluronan (HA), thereby forming HC-HA complexes (Colon et al., 2009, Jessen and Ødum, 2003, Mukhopadhyay et al., 2004). Due to this activity, TSG-6 has an important role in the stabilization and expansion of cumulus cell-oocyte complexes, a phenomenon vital for egg cell maturation; therefore female TSG-6 null mouse are sterile (Fulop et al., 1997, Fulop et al., 2003, Salustri et al., 1999). Additionally, TSG-6 protein is able to inhibit neutrophil migration via a competitive interaction involving cytokines and the Link domain of TSG-6 (Cao et al., 2004, Getting et al., 2002). The Link module interacts with the ECM on endothelial surfaces, and also with neutrophil-regulatory chemokines such as human CXCL8 (Dyer et al. 2016; Dyer et al. 2014). As elucidated by Dyer et al., TSG-6 competes with binding of chemokines to sulfated glycosaminoglycans of the ECM, thereby affecting the ratio of bound versus free cytokine. Thus, it is conceivable that the absence of TSG-6 could interrupt normal interactions that regulate the availability of free cytokine, thereby affecting neutrophil migration into wounds. A final consideration, although not a focus of our current manuscript, is published evidence that TSG-6 and HC-HA complexes can play a role in determining macrophage polarization (pro-inflammatory versus anti-inflammatory functions) in wounds (Ferrer et al., 2017, He et al., 2013, Mittal et al., 2016).

Given the ability of TSG-6 to regulate the behavior of neutrophils and macrophages, we postulated that TSG-6 may be important in cutaneous wound healing. Previous evidence for this idea was largely indirect. Multiple investigations in other inflammatory disease models, such as rheumatoid arthritis and inflammatory colitis, suggested that inflammation induces TSG-6 expression and that TSG-6 activity exerts anti-inflammatory effects (described more in the Discussion). In the skin, there were only a few previous studies on TSG-6. Of those, one demonstrated the presence of TSG-6 and heavy chains HC1 and HC2 in human scars and keloids (Tan et al., 2011). Another showed that intradermal implantation of mesenchymal stem cells (MSCs) in murine skin led to TSG-6 release; faster wound healing; and less fibrosis, macrophage activation, TNFα accumulation, and granulation tissue formation relative to wildtype (WT) control wounds (Qi et al., 2014). A third study investigated the effect of recombinant TSG-6 (rTSG-6) on rabbit ear wounds and showed that wounds injected with rTSG-6 healed with less hypertrophic scar than wounds receiving sham injections (Wang et al., 2015).. Thus, available studies suggested that TSG-6 may play an anti-inflammatory role in cutaneous wound healing. The aim of the current study was to test this hypothesis by examining how the loss of endogenous TSG-6 affects the normal course of wound repair in a knockout mouse model (TSG-6 null). The results might potentially provide better insight into the possibility of TSG-6 as a future therapy to enhance wound healing.

## RESULTS

### TSG-6 protein is present in unwounded skin, and its expression increases post wounding

To determine the normal expression levels of TSG-6 in intact and injured skin, Western analyses were performed on proteins from intact skin and full-thickness excisional wounds of WT mice (Figure 1a and b). TSG-6 is constitutively expressed at low levels in unwounded skin and increases by more than 4-fold as early as 12 h post-wounding. Expression peaks by 3 days, then drops but remains above baseline until at least day 10 (Figure 1b). This experiment showed that TSG-6 is upregulated after wounding, suggesting a possible functional role in wound repair.

**Figure 1.**
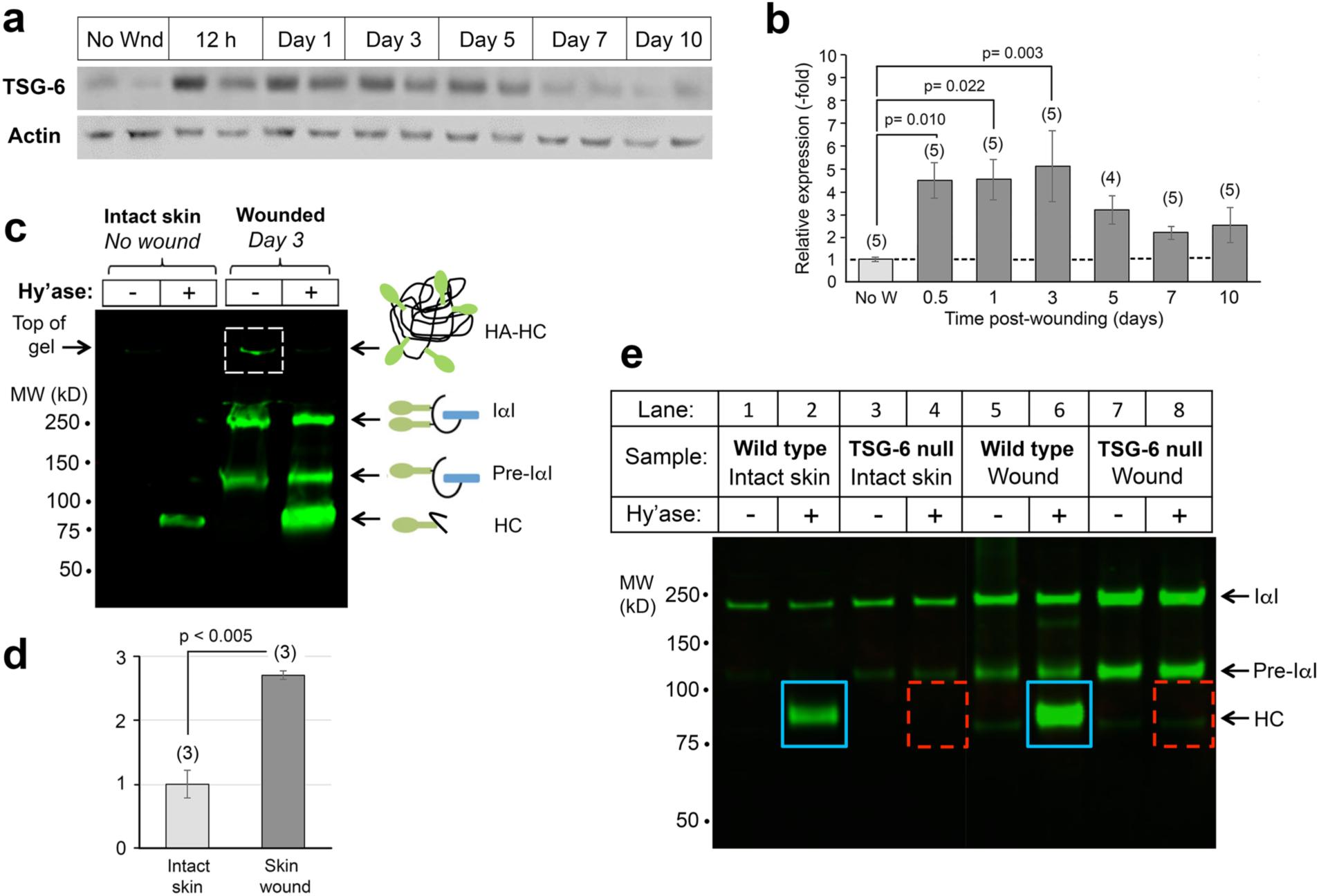
Expression of TSG-6 protein and its enzymatic product, HC-HA, in murine skin before and after wounding. (**a**), TSG-6 is expressed constitutively at low levels in uninjured skin, and highly induced by wounding. Western blots from wounds; two examples (different mice) are shown for each time point. (**b)**, Quantitation of TSG-6 protein expression from pooled samples, number of mice indicated (*n*); mean ± SEM; P*-*values from One-way Repeated Measures ANOVA. (**c**), Analysis of HC-HA in skin lysates on an acrylamide gel. HC proteins covalently bound to HA in HC-HA complexes are located at the top of the gel (boxed band, Lane 3). HC is released from complexes by degradation of HA with Streptomyces hyaluronidase (Hy’ase). Note that HC-HA levels are significantly higher in wounds (Lane 4) than in unwounded skin (Lane 2). (**d**), Quantitation of the HC band from 3 gel experiments; P-value, Student t test. (**e**) Gel analysis of HC-HA complexes in unwounded skin and in 24-hour excisional wounds, confirming that the amount of HC bound to HA is increased in wounds relative to intact skin (blue boxes). HA-bound HC is completely absent in TSG-6 null mouse skin, whether intact or wounded (dashed red boxes).

As mentioned earlier, TSG-6 catalyzes the covalent transfer of HCs from IαI and pre-αI to HA, forming HC-HA complexes. HC-HA is thought to stabilize HA rich extracellular matrices, making it more adhesive to inflammatory cells (Lauer et al., 2014, Rugg et al., 2005, Zhuo et al., 2006). To quantify the levels of HC-HA in unwounded and wounded WT skin, wound lysates were separated on PAGE gels as shown in Figure 1c; <u>details of this assay technique are given in Lauer 2015 and nicely summarized in Fig. 7 of Swaidani (Lauer et al., 2015, Swaidani et al., 2013).</u> Intact HC-HA complexes in the samples cannot enter the SDS-PAGE gel, and remain trapped in the well due to their large size (Figure 1c, lane 3, dashed box). However, with hyaluronidase treatment, HC-HA complexes are degraded and HC is released, appearing as a band at ∼85 kDa. <u>Larger bands in the gel (Pre-α−I, or I-α−I) correspond to bikunin proteins conjugated with one or two HC proteins, respectively</u>. The presence of a free HC band in the hyaluronidase-digested sample in Figure 1c, lane 2 shows that HC-HA is constitutively present in normal unwounded skin. As a result of wounding, the HC band intensity increases by ∼3 fold (Figure 1c, lane 4; quantified in Figure 1d). This change in HC-HA complexes mirrors the change in TSG-6 protein expression before and after wounding (Figure 1a and b).

### There are no enzymes other than TSG-6 that catalyze HC-HA transfer in murine skin

To determine whether TSG-6 protein is the sole molecule responsible for HC-HA complex formation in skin, we assayed HC-HA in extracts from the skin of TSG-6 null mice biopsied pre- and post-wounding. The HC band normally seen in unwounded WT samples (Figure 1e, lane 2) was completely absent in samples from unwounded TSG-6 null skin (Figure 1e, lane 4). At day 1 post-wounding, a time when HC-HA complexes are strongly induced in WT wounds (Figure 1e, lane 6), no HC band was detectable in TSG-6 null wounds (Figure 1e, lane 8). These data verify that loss of TSG-6 is responsible for the inability to form HC-HA complexes, and no other enzyme is expressed to compensate and transfer HC from IαI to HA.

### The absence of TSG-6 exerts major effects upon wound closure and neutrophil infiltration

Closure rate is an important indicator of proper wound healing. To observe the effects of loss of TSG-6 upon wound closure, we monitored the area of excisional wounds over time in WT and TSG-6 null mice. At all time points post-wounding, the relative area of macroscopically-observed wounds was significantly larger for TSG-6 null wounds than for WT wounds, demonstrating that loss of TSG-6 delays wound closure (<u>Figure 2a and 2b</u>). <u>Slower epithelial migration was one major component of this delayed wound closure, as confirmed by histological examination of biopsied wounds (Figure 2c and 2d)</u>.

**Figure 2.**
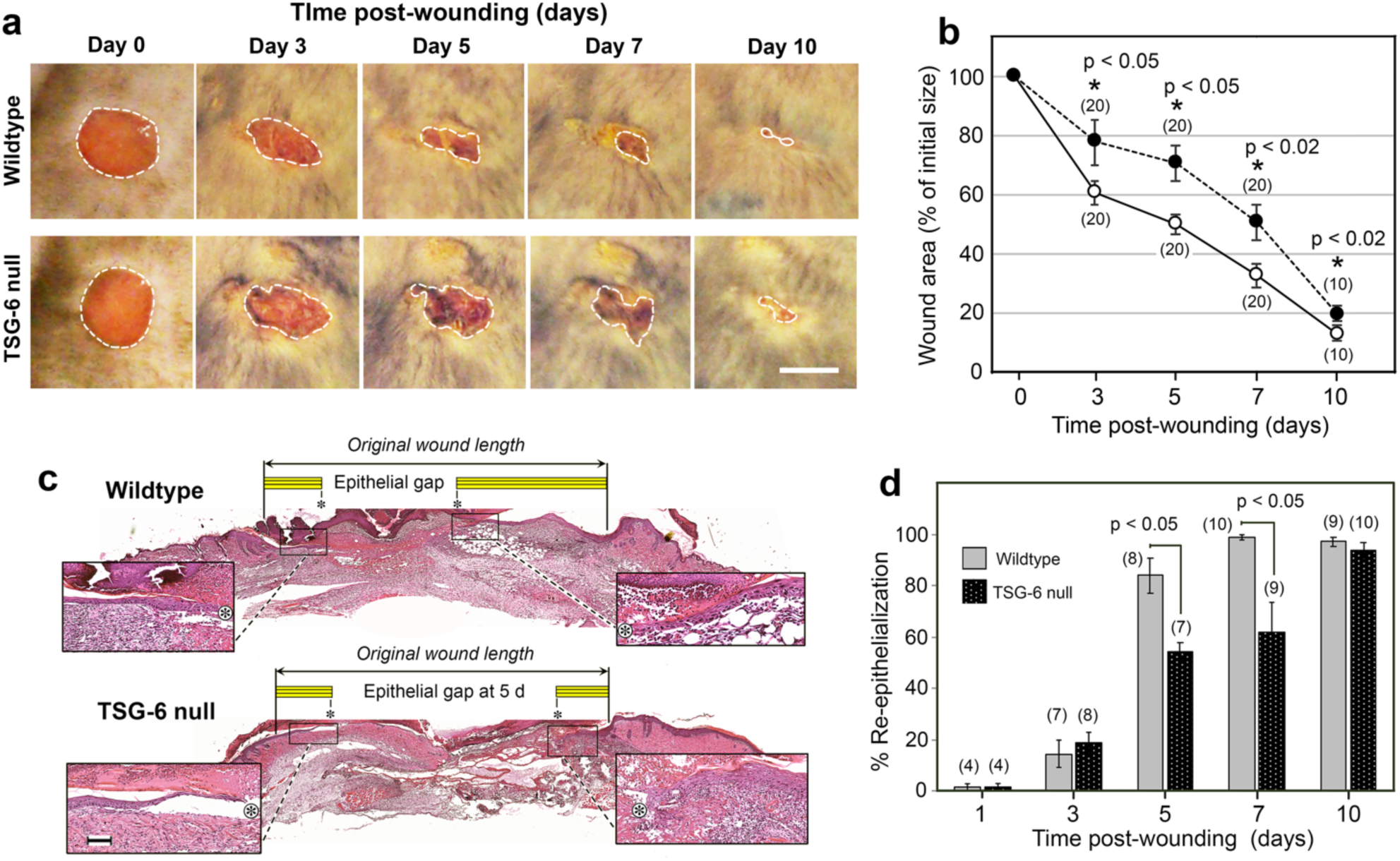
Wound closure is delayed in TSG-6 null mice. (**a)**, Circular 5-mm excisional wounds, created on the dorsum of mice (illustrated for one WT mouse and one TSG-6 null mouse were photographed every other day. *Scale bar*, 5 mm. (**b)**, Quantitation of macroscopic wound closure shows a significantly slower rate of closure in TSG-null mice, *dark circles*; mean ± SEM; (n), number of wounds; P-values from Student t-test (days 7 and 10) or Mann-Whitney Rank Sum test (days 3 and 5) after Shapiro-Wilk tests for normality. (**<u>c)</u>**<u>, Hematoxylin-and-eosin stained images of paraffin-fixed, full-thickness excisional skin wounds from a WT and a TSG-6 null mouse at 5 d post-wounding. Enlarged</u> *<u>insets</u>* <u>show the migrating epithelial tongues on either side of the wound;</u> *<u>Asterisk</u>*<u>, tip of epithelial tongue</u> *<u>Yellow bar</u>*<u>, length of migrating epithelium;</u> *<u>Scale bar</u>*<u>, 100 µm. (</u>**<u>d)</u>**<u>, Quantitation of histological wound closure from all wounds shows significant delay in reepithelialization in TSG-6 null mice relative to WT mice; (</u>*<u>n</u>*<u>), number of wounds; mean ± SEM; P-values from Mann-Whitney Rank Sum tests. Full-size images of re-epithelializing wounds, at all time points, can be viewed in</u> **<u>Supplementary Figure S1</u>**.

Another critical measure of proper wound healing is the orchestrated recruitment and clearance of inflammatory cells. Neutrophils are among the first to arrive at the site of injury. Interestingly, a biphasic pattern of neutrophil recruitment and/or clearance was observed in the presence *versus* absence of TSG-6 (and HC-HA) in cutaneous wounds. Sections from early wounds (12 h and 24 h) immunostained with the neutrophilic granule marker Ly6G showed significantly fewer neutrophils in the TSG-6 null mice compared to WT mice (<u>Figure 3a and 3c</u>). At 3 and 5 days post-wounding, neutrophil influx appeared relatively equal in WT and TSG-6 null, but by day 7 there were significantly more neutrophils in TSG-6 null wounds than in WT (Figure 3a and 3c). By day 10, neutrophils had returned to very low levels in the wound bed in both WT and null mice.

**Figure 3.**
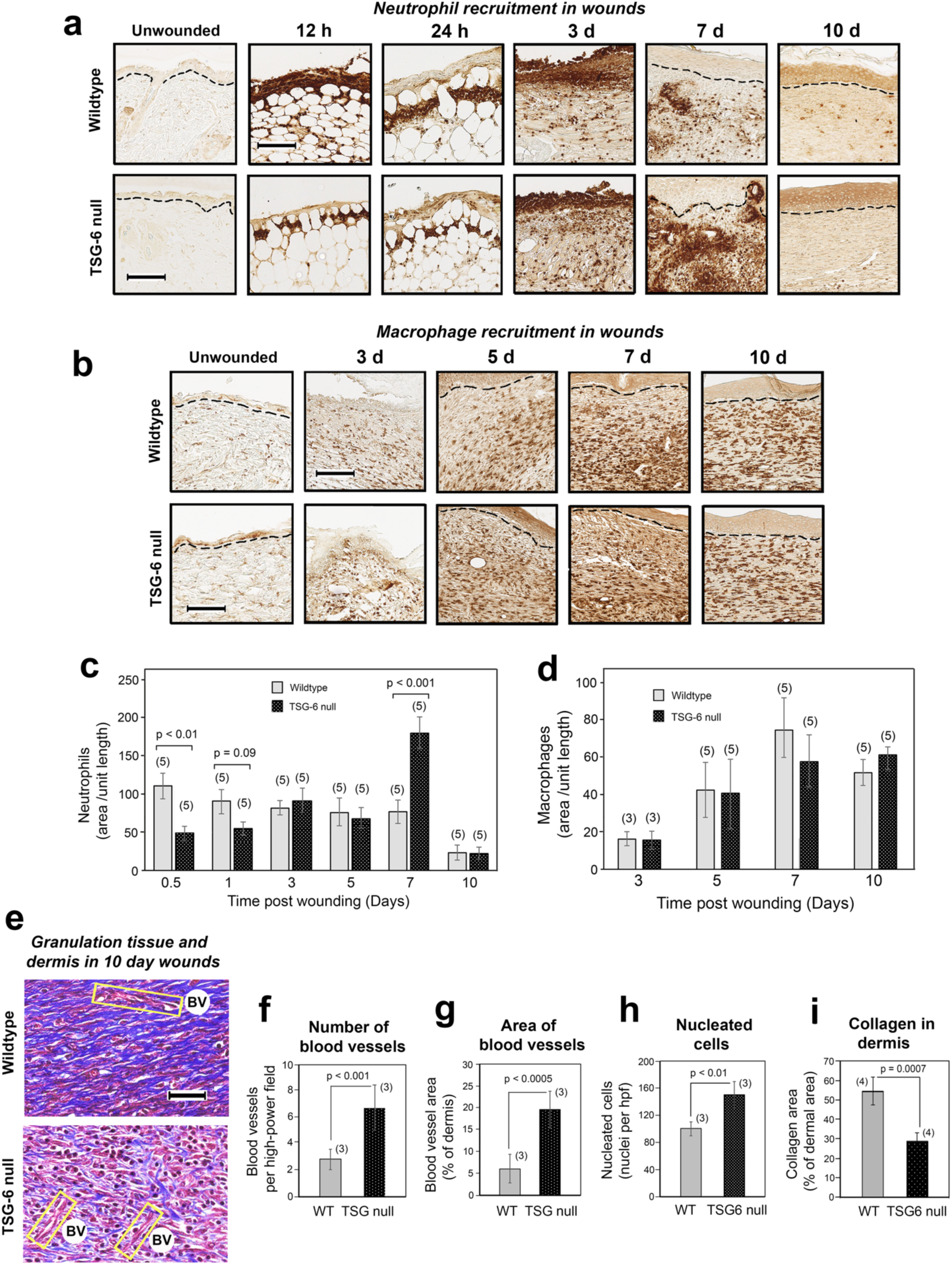
Time course survey of wound healing shows a biphasic abnormality in neutrophil recruitment, and a delay in resolution of the granulation phase, in TSG-6 null mice. (**a, c)**, Neutrophil influx is reduced at 12-24 h post-wounding, but increased at day 7, in TSG-6 null relative to WT wounds. **(a)**, Immunostains with anti -Ly6G antibody; scale bar, 100 µm. (**c)**, Quantification of neutrophils in the wound (see Methods); mean ± SEM; (n), number of animals; P values from Mann-Whitney Rank Sum tests. (**b, d**), Macrophage influx is equivalent in TSG-6 null and WT mice. (**b**), Immunostains with anti -F4/80 antibody; *scale bar*, 100 µm. (**d**), Quantification shows that macrophages are first detectable at day 3 and increase to a maximum level by day 7, with no difference between TSG-6 null and WT wounds. (**<u>e</u>**<u>), Masson-Trichrome staining of wound beds at 10 days post-injury for evaluation of blood vessels (BV; yellow boxes), tissue cellularity (red stain), and production of collagen (blue stain);</u> *<u>scale bar</u>*<u>, 50 µm.</u> **<u>(f - i)</u>**<u>, Triplicate or quadruplicate high-power fields (hpf) from stained wounds in 3 - 4 mice were analyzed using semi-quantitative image processing, to measure the number of blood vessels (</u>**<u>f</u>**<u>), blood vessel area (</u>**<u>g</u>**<u>), number of nucleated cells (</u>**<u>h</u>**<u>), and relative area of the dermis occupied by collagen (</u>**<u>i</u>**<u>). Full-size images of the wounds from which the enlarged panels in (</u>**<u>a</u>**<u>), (</u>**<u>b</u>**<u>), and (</u>**<u>e</u>**<u>) were selected can be viewed in</u> **<u>Supplementary Figures S2, S3</u>**<u>, and</u> **<u>S4</u>**<u>, respectively</u>.

To evaluate macrophages, we stained wound sections using the pan macrophage marker F4/80. No significant differences in the relative number of macrophages were observed between WT and TSG-6 null wounds, suggesting that the absence of TSG-6 and HC-HA had no significant effect upon total macrophage recruitment (<u>Figure 3b and 3d</u>).

<u>Late stages of wound healing (tissue-repair and remodeling) involve sequential creation of a vascularized provisional wound matrix (granulation tissue) that supports fibroblast neogenesis, followed by deposition of a well-organized collagenous matrix coinciding with involution of blood vessels. Masson-Trichrome staining of late-stage wounds at day 10 revealed clear-cut differences in the progression of tissue repair events in TSG-6 null</u> *<u>versus</u>* <u>WT mice (Fig. 3e-3i). At this time point post-wounding, WT wounds displayed extensive, well-organized collagen fibrils and very few blood vessels (as expected), whereas the TSG-6 null wounds showed persistence of many blood vessels and low amounts of collagen (Fig. 3e). Quantification of the stained sections revealed significantly more vasculature (Fig. 3f, g), more cellularity (Fig. 3h), and less collagen (Fig. 3i) in TSG-6 null wounds as compared to WT wounds, indicative of a defective maturation process in TSG-6 null healing wounds.</u>

### TSG-6 null mice develop an abnormally pro-inflammatory wound milieu

TSG-6 is known to have anti-inflammatory effects in other systems (Day and Milner 2018); thus the loss of TSG-6 might remove a brake on the expression of pro-inflammatory cytokines. To test this hypothesis, <u>we measured expression levels of TNFα in murine skin, both before and after wounding (Fig. 4). By Western analyses, TNFα protein levels were found to be highly elevated in TSG-6 null skin, even in the absence of injury (Fig. 4a, upper panel). While TNFα levels did increase after injury, especially at 12 h post-wounding, the relative difference between TSG-6 null and WT skin remained roughly the same (∼2-3 fold) throughout the first week of healing (Fig. 4b).</u>

**Figure 4.**
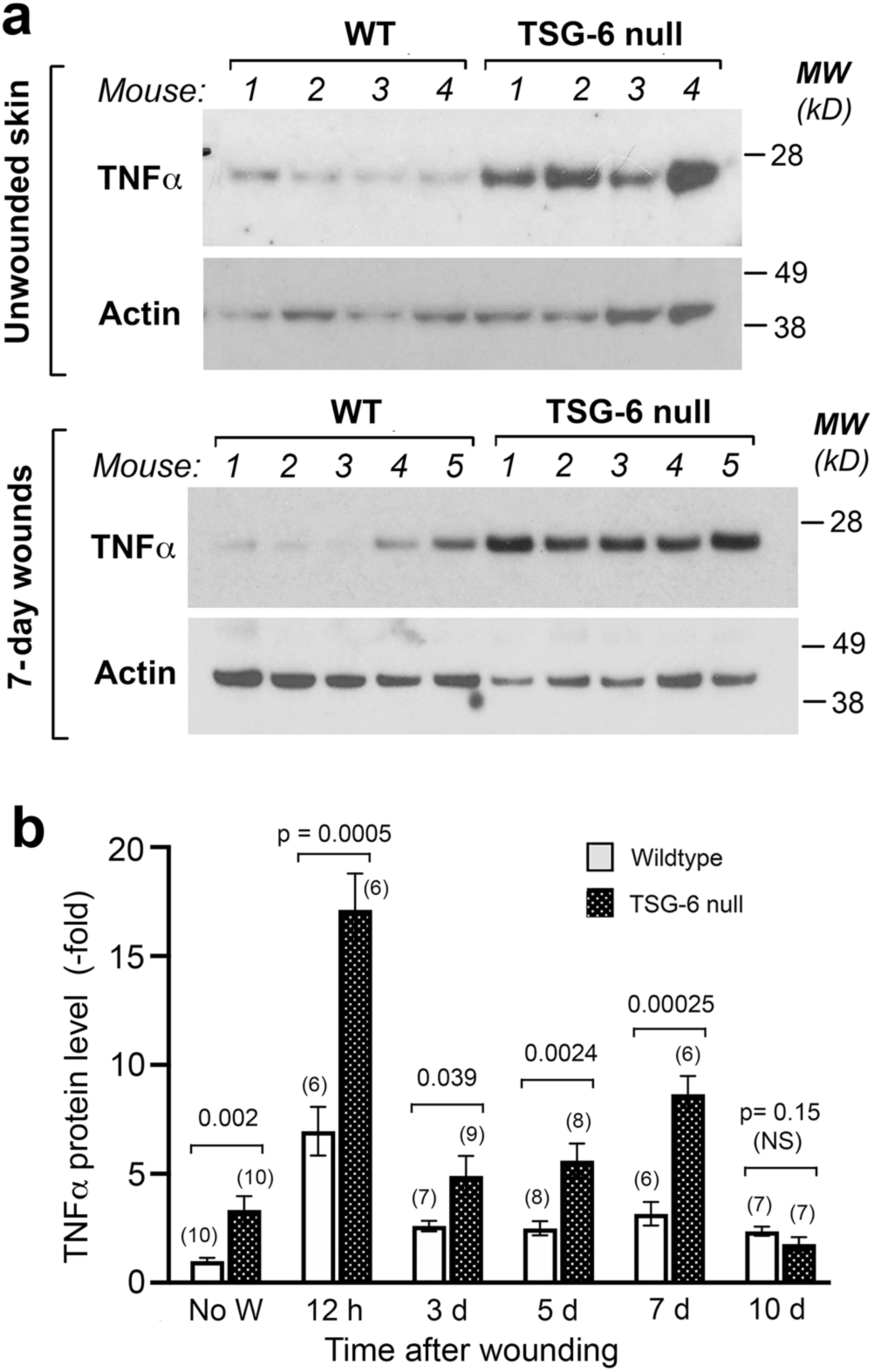
Loss of TSG-6 in skin upregulates the expression of the pro-inflammatory cytokine TNFα. (**<u>a</u>**<u>), Western analysis of unwounded skin (</u>*<u>upper panel</u>*<u>) and of 7-day healing wounds (</u>*<u>lower panel</u>*<u>), show higher TNFα protein expression in TSG-6 null mice as compared to WT mice. (</u>**<u>b</u>**<u>), Quantification of multiple Western blot experiments reveals significantly higher expression of TNFα in TSG-6 null skin, both before wounding and in the first 7 days after wounding; mean ± SEM; significance levels are indicated by P-values above the bars (Mann-Whitney Rank Sum test after Shapiro-Wilk tests for normality).</u>

<u>To examine possible involvement of other cytokines, lysates from wound biopsies at each of the relevant time points in our study were examined for expression levels of inflammation-related cytokines, including IL-6, KC, MCP-1, and IL-10, using a multiplexed ELISA assay (Supplementary Fig. 6). Although wounding-induced increases in expression were observed for each of these cytokines, no consistent differences between WT and TSG-6 null wounds were noted</u>.

### Reintroduction of recombinant TSG-6 into TSG-6 null wounds restores normal healing

To confirm that differences in wound closure between WT and TSG-6 null wounds are due to the loss of TSG-6, a protein rescue experiment was performed. Recombinant TSG-6 protein (rTSG-6) was injected into TSG-6 null wounds immediately after wounding (Day 0) and again at 4 days post-wounding (Figure 5a). After one injection, wound sizes trended slightly lower in TSG-6 null mice in wounds receiving the rTSG-6 injections (<u>Supplementary Fig. S5</u>). After the second injection, TSG-6 null wounds became significantly smaller than non-injected controls, and in fact equaled the size of WT wounds at day 7 (<u>Figure 5b</u> and c). Interestingly injection of rTSG-6 into WT wounds actually worsened wound closure compared to the WT vehicle control, indicating that TSG-6 may exert its closure effects within a critical concentration range. Overall, the findings of Fig. 5b and 5c suggest that the delay in wound closure in TSG-6 null wounds is attributable to loss of TSG-6.

**Figure 5.**
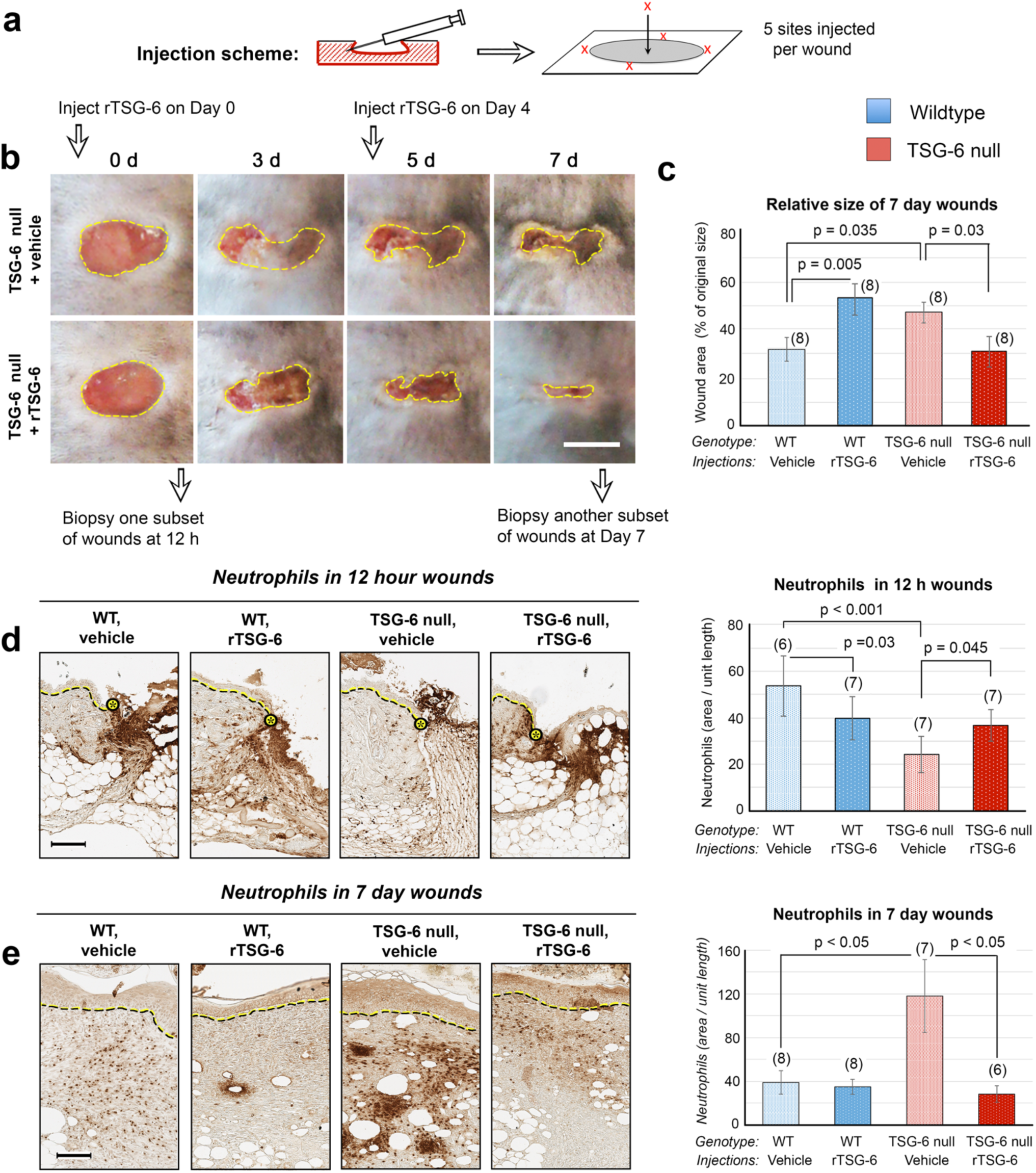
Reintroduction of recombinant TSG-6 (rTSG-6) into TSG-6 null wounds rescues the abnormal phenotype of delayed closure and abnormal neutrophil recruitment. (**a**), Schematic diagram showing the location of injections of rTSG-6 or vehicle into each wound. (**b**), Effect of rTSG-6 upon macroscopic wound closure. Photographs illustrate the effect in TSG-6 null mice when wounds were injected with rTSG-6, or with saline (vehicle) alone; *Scale bar*, 5 mm. (**c**), Quantification of wound closure, showing complete results at Day 7 in both WT and TSG-6 null mice, with either vehicle or rTSG-6 injections. (**d**), Neutrophil staining of wounds at 12 h; the delay in neutrophil recruitment in TSG-6 null wounds is partially rescued by rTSG-6 injections. *Dotted lines*, epidermis at wound edge. *Scale bar*, 100 µm. (**e**) Neutrophil staining of Day 7 wounds shows that the exacerbated neutrophil recruitment is completely reversed by rTSG-6 treatment. *Scale bar*, 100 µm. <u>For all graphs, bars show mean ± SEM. Statistical analysis (P-values) were performed using Two Way ANOVA with Bonferroni correction (graphs in c, d), or with Kruskal-Wallis One Way ANOVA (graph in e). A graph with closure data from all time points post-wounding is provided in</u> **<u>Supplementary Figure S5</u>**.

To confirm that differences in neutrophil recruitment between WT and TSG-6 null wounds are due to loss of TSG-6, another TSG-6 reintroduction experiment was performed. The experimental scheme was similar to that in Figure 5a except that wounds were harvested at 12 h or day 7 and histological sections were immunostained for Ly6G. Numbers of neutrophils at the wound site at 12 h (which were reduced in TSG-6 null mice relative to WT) were significantly increased after one injection of rTSG-6, becoming more similar to WT (<u>Figure 5d</u>). Likewise, the number of neutrophils in day 7 wounds, ordinarily much higher in TSG-6 null wounds, was completely normalized to WT levels in mice receiving two rTSG-6 injections (<u>Figure 5e</u>). Thus, the differential neutrophil recruitment observed in TSG-6 null wounds is reversible when TSG-6 is restored in the skin.

## DISCUSSION

This study describes abnormal wound healing responses in mice lacking TSG-6, an enzyme that catalyzes cross-linking of heavy chain proteins to hyaluronan, and regulates the local concentration of cytokines (soluble versus matrix-bound) in tissues. Our overall findings show that relative to WT controls, TSG-6 null wounds display (i) delayed wound closure, (ii) delayed neutrophil recruitment in early wounds, (iii) exacerbated neutrophil recruitment in late wounds, and (iv) increased levels of TNFα in normal skin and in the wound bed. Further, we demonstrated that reintroduction of recombinant TSG-6 can reverse both the delayed wound closure and the abnormal neutrophil recruitment.

Our study was motivated by previous literature suggesting an important role for TSG-6 in regulating inflammation. In many cell and tissue types, TSG-6 expression is low or absent under normal conditions, but is stimulated by pro-inflammatory factors such as TNFα, interleukin-1 (IL-1), and lipopolysaccharide (LPS) (Milner and Day, 2003). Here we show that TSG-6 is constitutively expressed at low levels in unwounded mouse skin, in agreement with earlier findings in human skin (Tan et al., 2011) and is greatly increased after wounding. Similarly, HC-HA complexes are detectable in low amounts in normal skin and become significantly elevated after wounding. However, HC-HA is completely absent in the skin of TSG-6 null mice. Therefore, levels of TSG-6 appear to correlate with levels of its enzymatic product, HC-HA. Regarding wound inflammation, results reported after injection of rTSG-6 or MSCs (which produce TSG-6 along with many other factors) in certain inflammatory models such as murine experimental arthritis (Bardos et al., 2001), myocardial infarction (Lee et al., 2009), DSS-induced colitis (Sala et al., 2015), and rabbit ear wounds (Wang et al., 2015), indicate that TSG-6 might have an anti-inflammatory role, resulting in decreased disease activity and/or improved recovery. In accordance with these studies, we found that absence of endogenous TSG-6 is deleterious to wound healing, whereas restoration of the protein through direct dermal injection of rTSG-6 into TSG-6 null mouse wounds tended to normalize wound healing, at least in terms of the parameters evaluated here.

Our study uncovered interesting defects in the recruitment of neutrophils in the wound bed of TSG-6 null mice. As a brief review, wounding initiates the release of multiple proinflammatory factors that stimulate local recruitment of neutrophils and monocytes into the wound bed (Martin and Leibovich, 2005, Singer and Clark, 1999, Wang, 2018). Neutrophil recruitment starts as early as 4 h post-wounding and plateaus around 3 days (Kim et al., 2008). When TSG-6 has been studied in other model systems, the effects on neutrophils appeared consistent with an inhibitory effect. Thus, injection of rTSG-6 protein into inflamed tissue tended to decrease the inflammatory response, including reductions in pro-inflammatory cytokines and neutrophil infiltration (Beltran et al., 2015, Kim et al., 2014, Oh et al., 2010). Absence of TSG-6, on the other hand, led to increased neutrophil infiltration (Szanto et al., 2004). In our TSG-6 null wound model, the neutrophil recruitment response appears to be biphasic. At early times (12 and 24 h post-wounding), neutrophils are significantly lower in TSG-6 null than in WT wounds. The explanation for this is only speculative, but one idea could be an interaction between neutrophils and HC-HA on the endothelial glycocalyx. This is based on the known fact that the luminal side (endothelium) of blood vessels is lined with an HA rich glycocalyx (Reitsma et al., 2007, Shakya et al., 2015), which might under wound conditions become decorated with HC, forming HC-HA complexes. Leukocytes were shown to bind more avidly to HC-HA complexes than to non-complexed HA (Lauer et al., 2014, Zhuo et al., 2006); therefore HC-HA might facilitate neutrophil attachment within small vessels in the wound bed in WT wounds. In that case, the absence of HC-HA in TSG-6 null wounds might delay neutrophil recruitment in early wounds.

In late wounds, the mechanisms of neutrophil recruitment are apparently different than in early wounds, as there are more neutrophils in the late TSG-6 null wounds than in corresponding WT wounds. <u>One possible explanation is that absence of TSG-6 results in increased expression of inflammatory cytokines during later stages of wound repair. However, in testing that hypothesis, our measurements of whole-wound lysates (even when using sensitive Milliplex protein assays) were unable to demonstrate any meaningful differences (Supplementary Fig 6). Another hypothetical reason for differences in the late WT versus TSG-6 null wounds could be that one or more pro-inflammatory cytokines such as KC (murine IL-8, a neutrophil chemoattractant) are more available in TSG-6 null wounds than in WT wounds. TSG-6 is known to bind to many of these cytokines, and to compete with cytokine binding to ECM proteoglycans in assays using purified reagents measured by surface plasmon resonance (Dyer et al., 2016, Dyer et al., 2014). Thus it remains possible that TSG-6 is, in fact, crucial for regulating the availability of ‘free’ cytokine binding to their cognate receptors on cell membranes, but our method (examination of total extracts from whole skin lysates) was insufficient to reliably detect differences between bound and free cytokines.</u> Regardless of the nature of the neutrophil recruitment mechanism, excessive neutrophil activity has been tied to delayed wound healing (Kim et al., 2008, Wilgus et al., 2013), and therefore likely contributes to the delayed wound healing observed in TSG-6 null mice.

<u>Interestingly in our study, TNFα expression was found to be consistently higher in TSG-6 null skin, not only after injury (12h and out to 7 days), but also in unwounded skin. The latter finding is especially interesting because TSG-6 (TNF-stimulated gene 6) was first discovered as a gene whose expression is inducible by TNFα (Lee et al., 1990). The fact that elevated levels of TNFα are observed when TSG-6 is missing may indicate the existence of a previously-unrecognized feedback loop in which TNFα gene expression is normally suppressed by TSG-6, but becomes de-repressed in the absence of TSG-6</u>. More detailed studies are planned to investigate this possibility.

A particular strength of our study is that TSG-6 restoration (rTSG-6 injection into TSG-6 deficient wounds) was shown to normalize wound closure and neutrophil recruitment. These results unambiguously establish that the defects observed in TSG-6 null mice result from loss of TSG-6 protein. Interestingly, while reintroduction of rTSG-6 into TSG-6 null wounds reversed the wound delay and neutrophil recruitment defects, addition of rTSG-6 into normal WT wounds worsened the closure rate. Apparently, tight regulation of TSG-6 is critical for proper wound healing, and either excessive or insufficient amounts can be deleterious.

In summary, loss of TSG-6 leads to abnormal pro-inflammatory changes and is detrimental to wound healing, whereas a regulated amount of TSG-6 is crucial for a normal inflammatory response and proper wound closure. While more studies are needed to fully elucidate the mechanisms behind the phenotypes observed, our study provides important groundwork for understanding the role of endogenous TSG-6 in cutaneous wound healing.

## MATERIALS AND METHODS

### Animals

C57BL/6J mice were obtained from JAX Laboratories (Bar Harbor, ME). TSG-6 null mice, whose generation was described by Fülöp et al. (Fulop et al., 2003), was generously provided by Dr. Mark Aronica in our institute. Both male and female mice, in approximately equal numbers, were used in these experiments. Mice were maintained per guidelines of the American Association for the Accreditation of Laboratory Animal Care. All procedures were approved by the Cleveland Clinic Institutional Animal Care and Use Committee (IACUC).

### Wounding experiments

Mice (8–10 weeks old) were anesthetized via intraperitoneal Ketamine-Xylazine injection, and the upper back shaved with an electric razor. The next day, two full-thickness excisional wounds were made using 5 mm punch biopsies (Acuderm, Fort Lauderdale, FL) under anesthesia. To follow wound closure, mice were anesthetized with isofluorane and photographed from a fixed distance using a digital camera. Wound area was determined from the photos using standard imaging software. For histological examination, wound tissues were harvested at defined time points, fixed in Histochoice (Amresco, Solon, OH), and paraffin embedded. Other wound tissues were flash frozen in liquid nitrogen and stored at -80 °C for subsequent protein and RNA isolation.

### Western Blotting

Protein lysates were prepared from wound samples as described previously (Mack and Maytin 2010). Lysates were electrophoresed in NuPage 4-12 % Bis-Tris gels (Invitrogen, Carlsbad, CA), blotted onto PVDF membranes (Millipore, Burlington, MA), and probed overnight (4 °C) with one of the following primary antibodies: Anti-TSG-6 mouse monoclonal (1 μg/ml; Millipore); rat monoclonal TNFα (1 μg/ml; BioXCell, West Lebanon, NH); or rabbit anti-mouse Actin (1:1000; Santa Cruz Biotechnology, Dallas, TX). This was followed by 1.5 hour (room temperature) incubation with appropriate horseradish peroxidase (HRP)-conjugated secondary antibodies (1:10,000; Jackson ImmunoResearch, West Grove, PA). Chemiluminescent detection was performed using Amersham™ ECL™, or ECL™ Prime reagent kits (GE Healthcare, Chicago, IL). Western blots were first probed with the antibody of interest, then stripped and reprobed for actin as a loading control. Protein bands were quantified using IPLab Spectrum imaging software (Scanalytics, Rockville, MD). Band intensities were first corrected by local background subtraction, then normalized to actin, and finally normalized to the band intensity of the protein in unwounded wildtype skin (Fig 1b and 4b).

### Evaluation of HC modification of hyaluronan (HC-HA analysis)

HC-HA analyses were performed as described (Lauer et al. 2015). Briefly, skin tissues were collected, weighed and minced in PBS at 1 μl volume per 0.33 mg of tissue. *Streptomyces* hyaluronidase (Millipore) was added (20 TRU/ml for extraction of HCs, or equal volumes of PBS for No-hyaluronidase controls), incubated for 30 min on wet ice followed by 30 min at 37°C. After centrifugation (13,000 rpm), supernatants were collected, electrophoresed, and blotted. Blots were incubated with anti-IαI rabbit polyclonal Dako antibody (1:8000; Agilent Technologies, Santa Clara, CA), followed by IRDye® 800CW donkey anti-rabbit secondary (1:15,000; LI-COR Biosciences, Licoln, NE). Blots were imaged on an Odyssey infrared imaging system (LI-COR Biosciences).

### Histochemical staining methods

Paraffin-fixed wound sections were microtome-sectioned, and stained with hematoxylin/eosin, or with Masson Trichrome stains. <u>Further details are in the Supplementary Figures</u>.

### Immunostaining for neutrophils and macrophages

Paraffin embedded tissues were cut (5-micron sections), rehydrated, blocked with 3% normal goat serum for 30 min. Sections were incubated overnight (4 °C) with the respective primary antibodies: Anti-Ly6G rat monoclonal (1:100; Affymetrix, Santa Clara, CA) for neutrophils; Anti F4/80 rat monoclonal (1:100; Bio-Rad, Hercules, CA) for macrophages. A biotinylated goat anti-rat biotin secondary (1:300) was followed by Vectastain ABC reagent (Vector Laboratories, Burlingame, CA) for 30 minutes each. After detection using a DAB peroxidase substrate kit, mounted slides were scanned using an Aperio AT2 slide-scanner (Leica Biosystems, Buffalo Grove, IL). Areas positively stained for neutrophils or macrophages were evaluated using IPLab software (Scanalytics).

### Quantification of histomorphologic parameters

Detailed methods for quantifying wound parameters such as wound length, staining intensity, or cell numbers using image processing of digital images of our microscopic sections are described in Supplementary Figures S1-S4.

### Recombinant TSG-6 injection

Mice were shaved, anesthetized, and two full thickness excisional wounds made using 5 mm punch biopsies. The edges and center of each wound was injected using an insulin syringe, with 2 µg of recombinant human TSG-6 (rTSG-6, R&D Systems, Minneapolis, MN) in 100 µl PBS (or with PBS alone) immediately after wounding and at Day 4 post wounding. Wounds were photographed to evaluate closure. Some wounds were collected at 12 h or Day 7 post wounding to evaluate neutrophil and macrophage infiltration.=

### Statistical analysis

Data were first tested for normality using a Shapiro-Wilk test. Normally distributed data were analyzed using a two-tailed Student’s t-test or Two-way Analysis of Variance, whereas data with a nonparametric distribution were analyzed using the Mann-Whitney Rank Sum Test. A *p*-value < 0.05 was considered statistically significant.

## DATA AVAILABILITY STATEMENT

No datasets were generated or analyzed during the current study.

## ACKNOWLEDGEMENTS

We thank Katie Hardin for her work on image analysis of neutrophils, Dr. Mark Aronica for sharing the C57BL/6 TSG-6 null mice, and Cassandra Singer for help with the Millipore/Sigma cytokine assay protocol. We acknowledge the teaching and contributions of the late Mark Lauer, Ph.D. and of Dr. Vincent Hascall, a champion of all things related to hyaluronan.

This work was supported by grant P01 HL107147 from NIH/NHLBI.

## CReditT STATEMENT

*Shakya, Sajina:* Conceptualization, Formal analysis, Investigation, Methodology, Writing (lead).

*Mack, Judith:* Conceptualization, Investigation, Methodology, Project administration, Supervision (lead), Writing (supporting).

*Alipour, Minou:* Formal analysis, Investigation, Methodology.

*Maytin Edward:* Conceptualization, Formal analysis, Funding acquisition (lead), Methodology, Resources (lead), Writing (supporting).

