## Supplementary material for "Cutaneous Wounds in Mice Lacking Tumor Necrosis Factor-stimulated Gene-6 Exhibit Delayed Closure and an Abnormal Inflammatory Response": Supplem Figures S1-S6

For an article in *J Invest Dermatol* by

All figures show comparisons between wildtype mice and TSG-6 null mice.

TABLE OF CONTENTS PAGE

**Supplementary Figure S1:**
Time course of re-epithelialization of wounds 2-4

**Supplementary Figure S2:**
Time course of Neutrophil recruitment in excisional skin wounds 5-8

**Supplementary Figure S3:**
Time course of Macrophage recruitment in excisional skin wounds 9-11

**Supplementary Figure S4:**
Transition from granulation stage to tissue maturation stage in late skin wounds 12-14

**Supplementary Figure S5:**
Rescue of the delayed closure phenotype in TSG-6 null mice by injection of rTSG-6 protein 15

**Supplementary Figure S6:**
Analysis of cytokines in skin wounds of TSG-6 null and wildtype mice 16

**Supplementary Figure S1**.

**Time course of re-epithelialization of wounds in wildtype mice and TSG-6 null mice.**

**Figure legend:**

Hematoxylin-eosin stained images of representative paraffin-fixed, full-thickness excisional skin wounds from wildtype (**a, c, e, g**) or TSG-6 null (**b, d, f, h**) mice. Wounds were harvested at various times post-injury, as indicated above each panel. Enlarged *insets* illustrate the migrating epithelial tongue found on each side of the wound. *Asterisks* indicate the distal edge of the epithelial tongue. Lengths of the re-epithelialized tissue are indicated by horizontal yellow bars. The width of the original wound is shown by arrow-tipped horizontal lines between the vertical lines at the edges of intact, follicle-bearing skin.
*Scale bar*, 100 μm.

**Experimental details:**

Mice (WT, or TSG-6 null, male and female, 8–10 weeks of age) were anesthetized with an intraperitoneal injection of Ketamine-Xylazine and fur shaved from the upper back with an electric razor. After letting the skin recover overnight, two full-thickness excisional wounds were made using 5 mm punch biopsies (Acuderm, Fort Lauderdale, FL), under anesthesia. Wounds were harvested at 12 h, and at days 1, 3, 5, 7 and 10 post wounding under deep anesthesia, followed by euthanasia. Histochoice-fixed, paraffin embedded wound tissues were cut into 5-micron sections using a standard microtome. After rehydration, sections were stained for Hematoxylin and Eosin (H&E) using standard procedures. Mounted slides were then scanned using an Aperio AT2 slide-scanner (Leica Biosystems, Buffalo Grove, IL). Scanned images were viewed and analyzed for histological wound closure using Aperio ImageScope (Leica Biosystems). Percentage re-epithelialization (or epidermal gap) was determined by measuring (A) the length of the epidermal tongue at both wound edges, and (B) the total wound length from the wound edge-to-edge. Percent epithelial closure was then expressed as the ratio A/B (see Fig. 2d).

Supplementary Figure S1, panels a-d

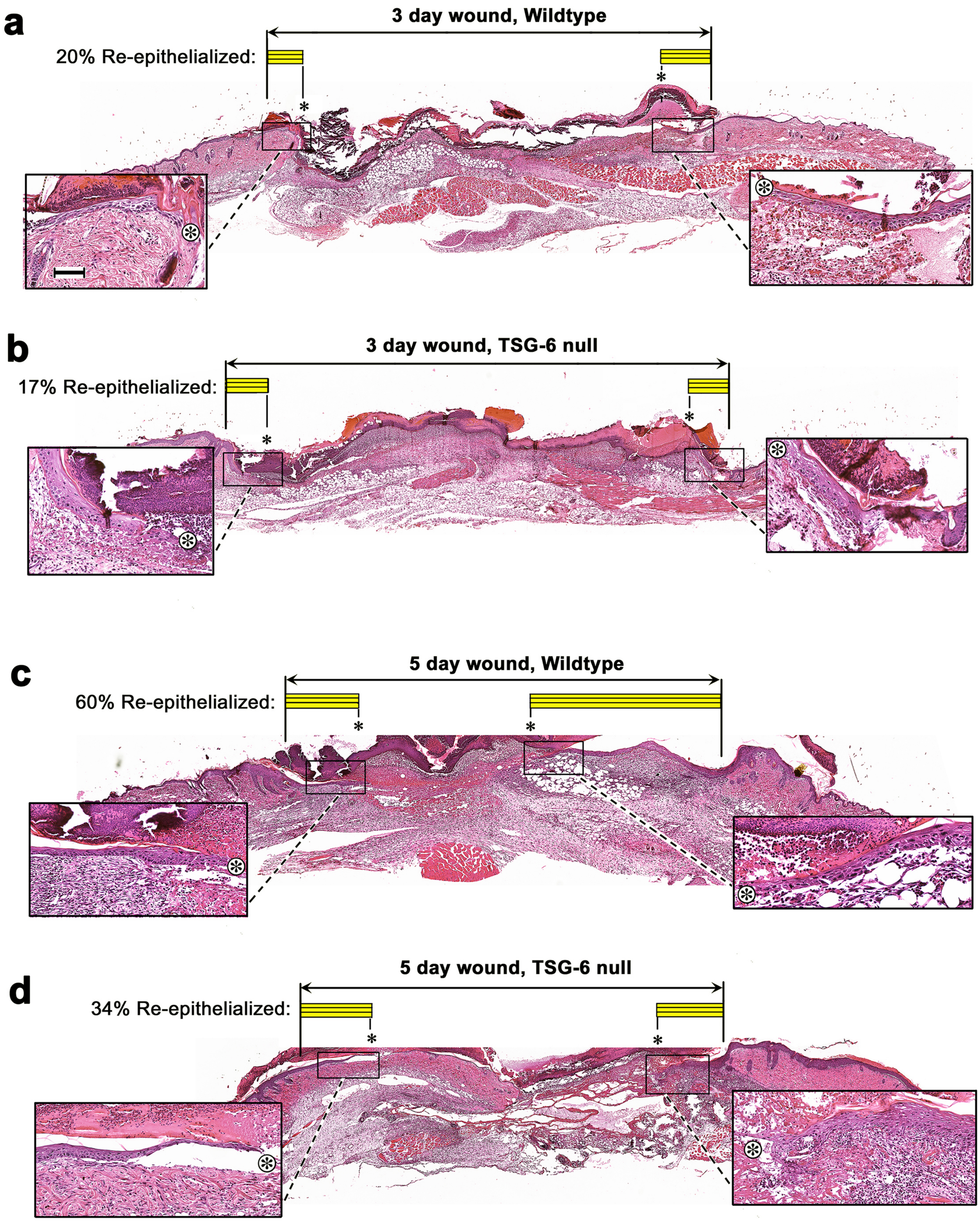

Supplementary Figure S1, panels e-h

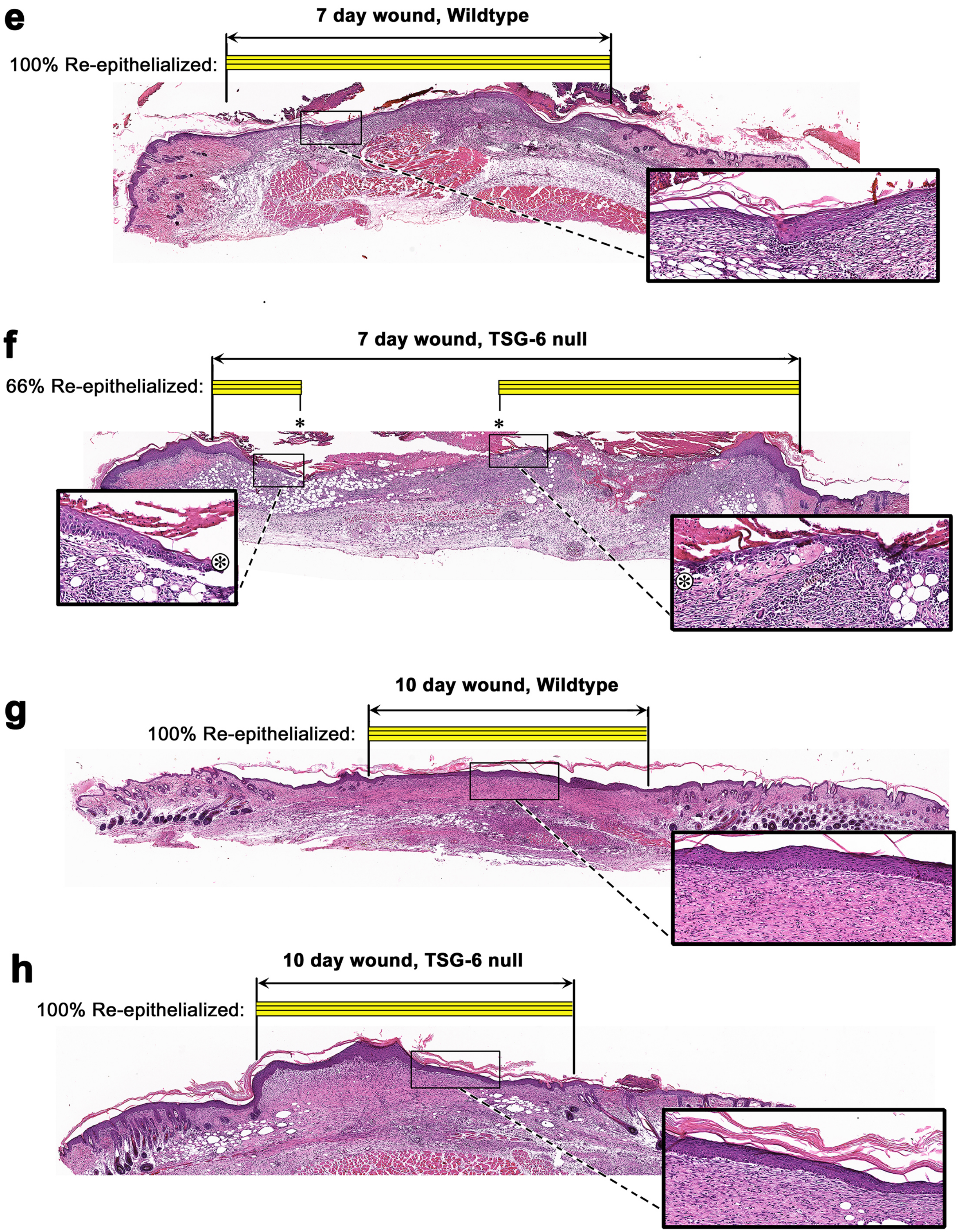

### Supplementary Figure S2.

**Neutrophil recruitment in excisional skin wounds in wildtype and TSG-6 null mice.**

**Figure legend:**

Skin wounds were harvested at various times post-injury, then were paraffin-fixed and immunostained with anti-Ly6G antibody for neutrophils and a brown peroxidase substrate for detection. Enlarged *insets* illustrate the appearance of neutrophils at the wound edge (*left panels*), and in the wound bed (*right panels*). The horizontal black line above each panel denotes the original wound edges identified by hair follicles in the intact perilesional skin. *Scale bar*, 100 μm.

This figure illustrates the type of data used to generate our semiquantitative analysis of neutrophils, as described in more detail below, and reported in Fig. 3 and Fig. 5 of the manuscript. Note that relatively *fewer* neutrophils are found at 12 h and 24 h in TSG-6 null wounds as compared to wildtype control wounds. Conversely, at 7 d post-wounding relatively *more* neutrophils are present in the TSG-6 null wounds than in wildtype wounds.

**Experimental details:**

Mice received 5 mm full-thickness wounds which were harvested at either 12 h, 24 h, 3 d, 5d, 7d, or 10d post-wounding, as previously described. Histochoice-fixed, paraffin embedded wound tissues were cut into 5-micron sections, rehydrated, blocked with 3% normal goat serum for 30 min at room temperature, then incubated at 4 °C overnight with Anti-Ly6G rat monoclonal (1:100; Affymetrix, Santa Clara, CA) which detects neutrophils. A biotinylated goat anti-rat biotin secondary (1:300) was followed by Vectastain ABC reagent (Vector Laboratories, Burlingame, CA), for 30 min each. Detection was performed using a DAB peroxidase substrate kit, followed by mounting in VectaMount mounting medium (Vector Labs). Slides were scanned using an Aperio AT2 slide-scanner (Leica Biosystems). To generate the quantitative data shown in see Figures 3 and 5, IPLab Spectrum imaging software (Scanalytics) was used to quantify the positive staining for neutrophils in the wound bed. A common threshold was set to encompass the majority of pixels with dark brown DAB positive staining; this positively-stained area was then expressed per unit wound length to provide normalization between samples.

Supplementary Figure S2, panels a-d

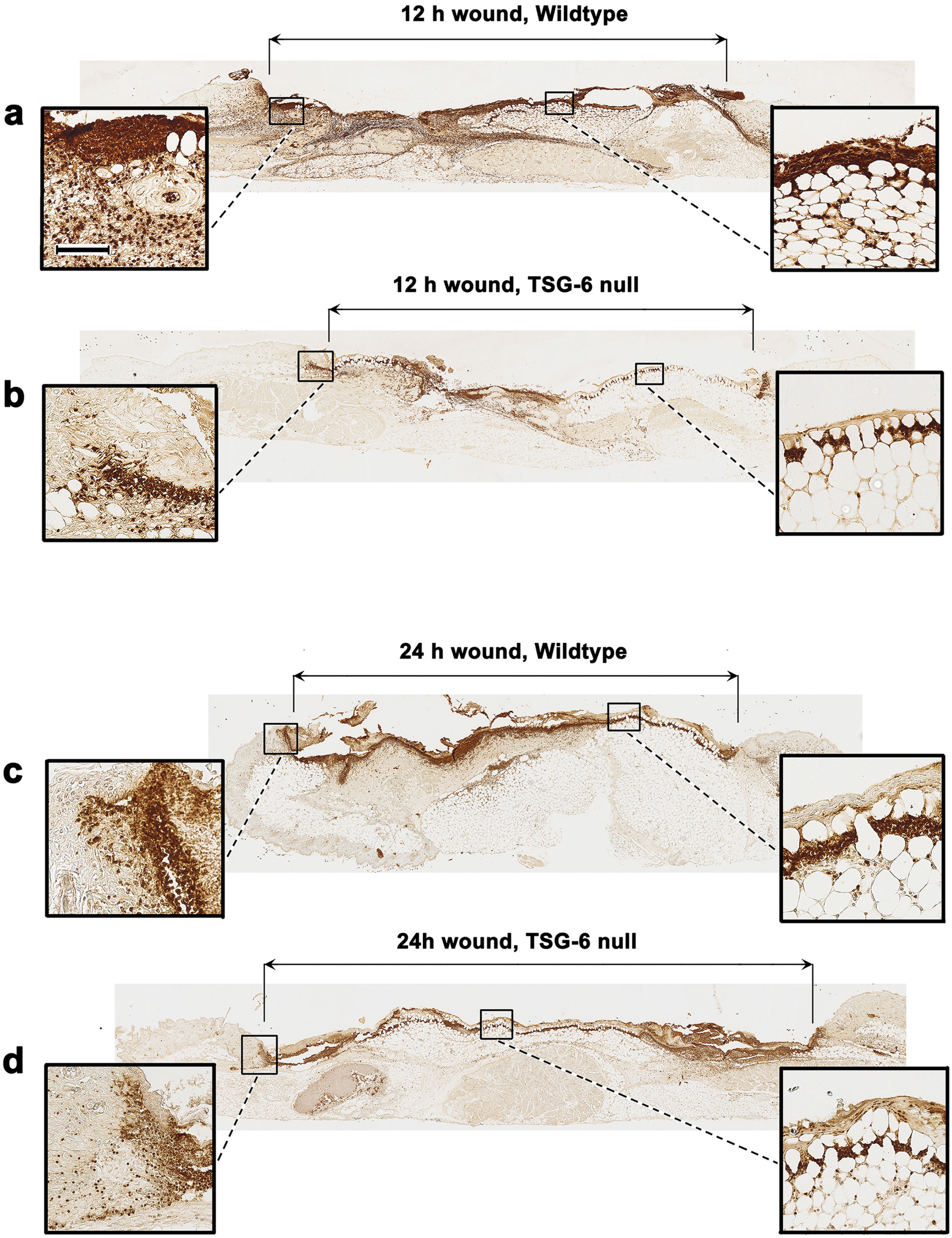

Supplementary Figure S2, panels e-h

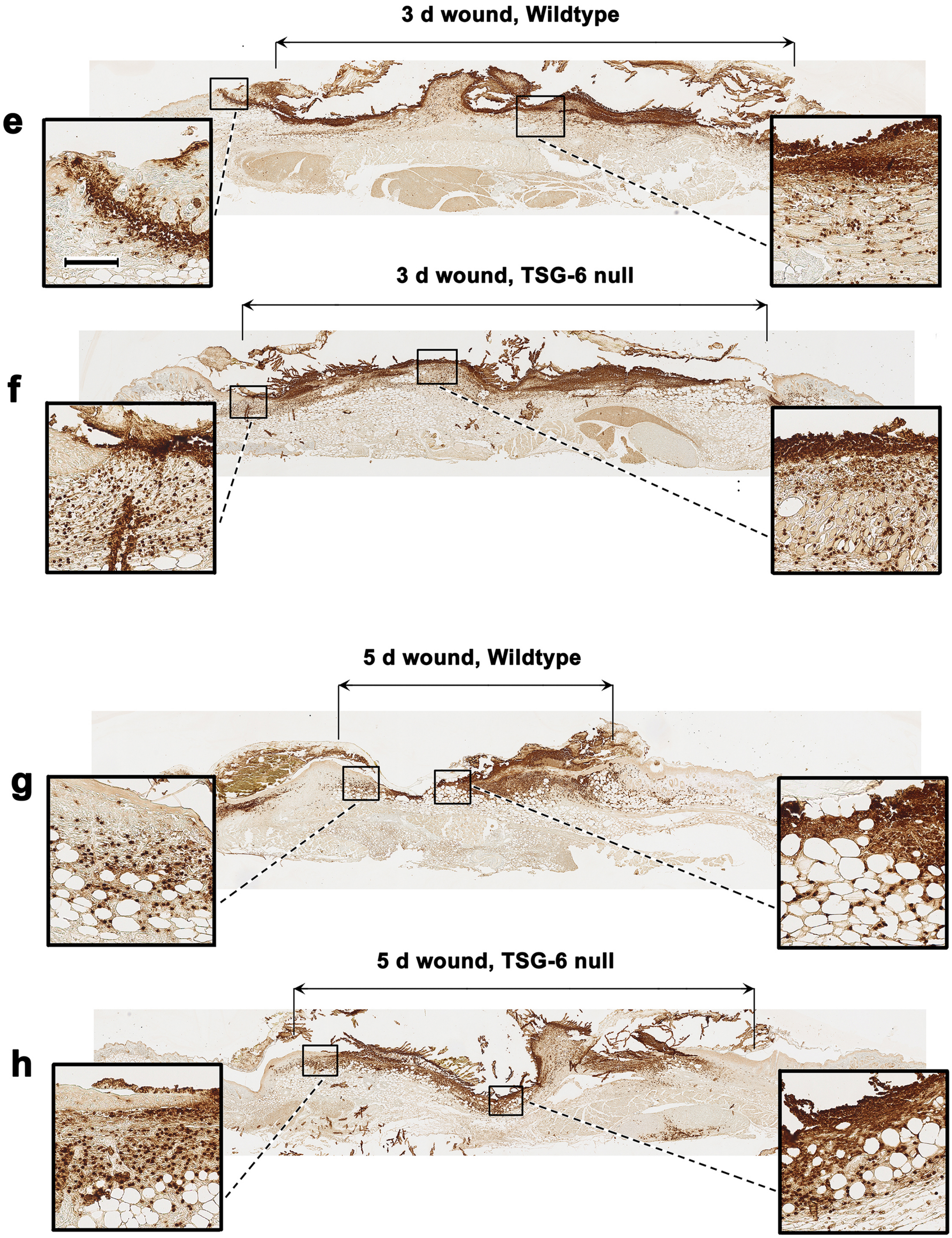

Supplementary Figure S2, panels i-L

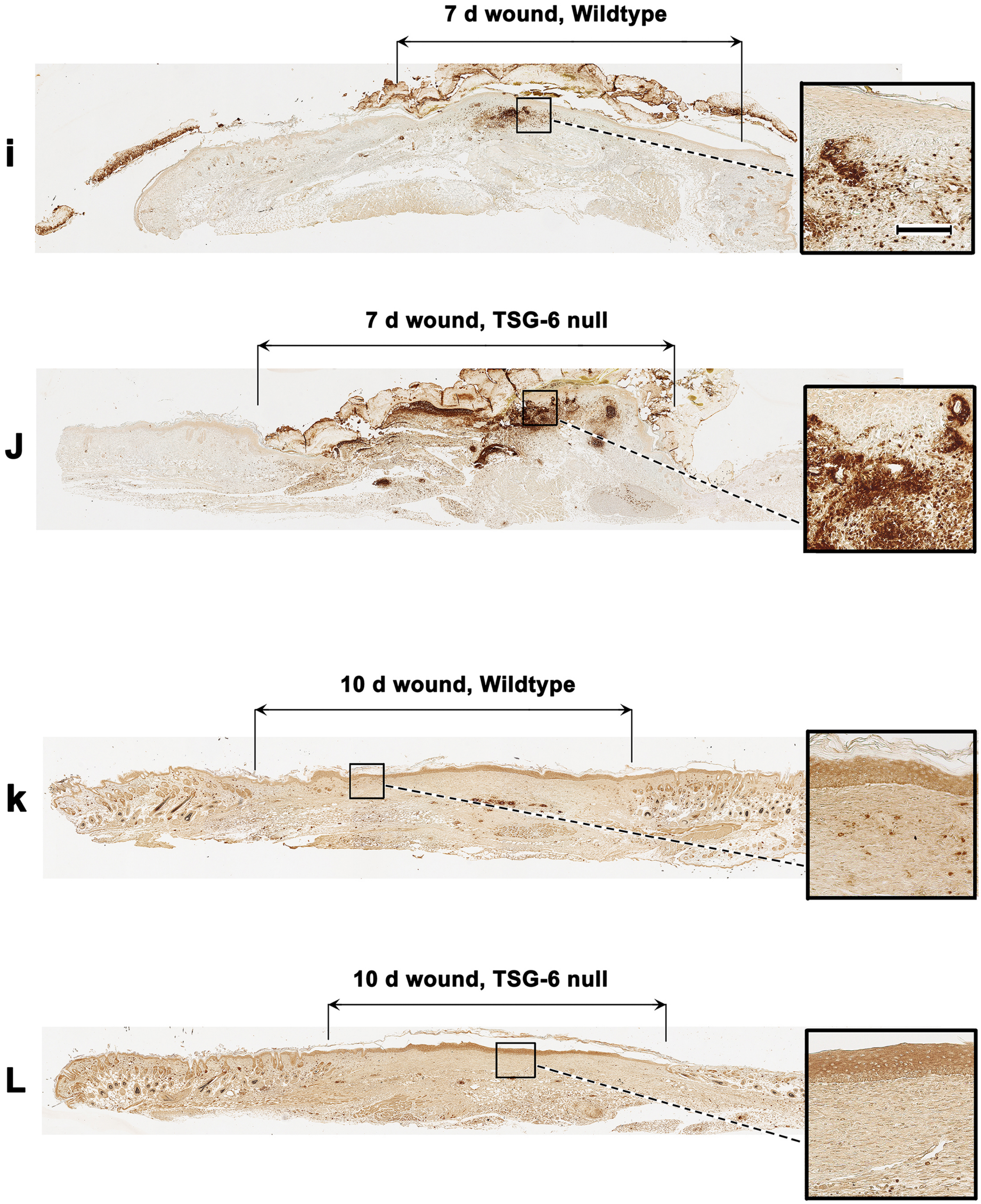

**Supplementary Figure S3**.

**Macrophage recruitment in excisional skin wounds in wildtype and TSG-6 null mice.**

**Figure legend:**

Skin wounds were harvested at various times post-injury, then were paraffin-fixed and immunostained with an anti-F4/80 antibody for macrophages followed by a brown peroxidase substrate for detection. Enlarged *insets* illustrate the appearance of the macrophages in the wound bed. The horizontal black line above each panel indicates the original wound edges as identified by hair follicles in the intact perilesional skin. *Scale bar*, 100 µm.

This figure illustrates the type of data used to generate the semiquantitative analysis of macrophage influx, as described in more detail below and reported in Fig. 3 of the manuscript. Unlike for neutrophils, macrophages show no relative differences in number in the TSG-6 null wounds relative to wildtype wounds. Negligible F4/80 stained macrophages were detected in 12 h or 24 h wounds (*data not shown*).

**Experimental details:**

Mice received 5 mm full-thickness wounds which were harvested at either 12 h, 24 h, 3 d, 5d, 7d, or 10d post-wounding, as previously described. Histochoice-fixed, paraffin embedded wound tissues were cut into 5-micron sections, rehydrated, blocked with 3% normal goat serum for 30 min at room temperature, then incubated at 4 °C overnight with anti-F4/80 rat monoclonal (1:100; Bio-Rad, Hercules, CA) to detect macrophages. A biotinylated goat anti-rat biotin secondary (1:300) was followed by Vectastain ABC reagent (Vector Laboratories, Burlingame, CA), for 30 min each. Detection was with a DAB peroxidase substrate kit, followed by mounting in VectaMount mounting medium (Vector Labs). Slides were scanned using an Aperio AT2 slide-scanner (Leica Biosystems). To generate the quantitative data shown in see Fig. 3, IPLab Spectrum imaging software (Scanalytics) was used to quantify the positive staining for neutrophils in the wound bed. A common threshold was set to encompass the majority of pixels with dark brown DAB positive staining; this positively-stained area was then expressed per unit wound length to provide normalization between samples.

Supplementary Figure S3, panels a-d

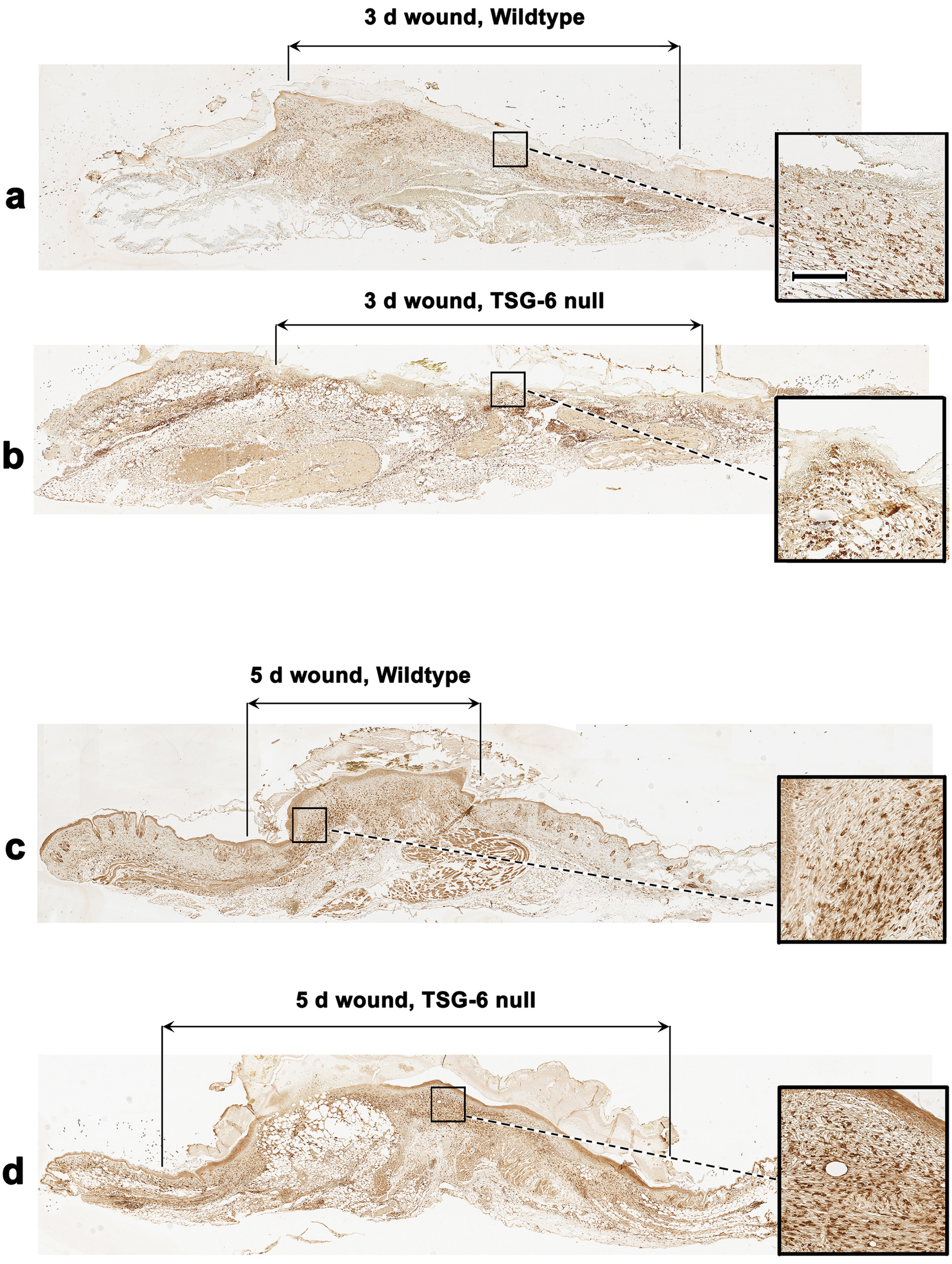

Supplementary Figure S3, panels e-h

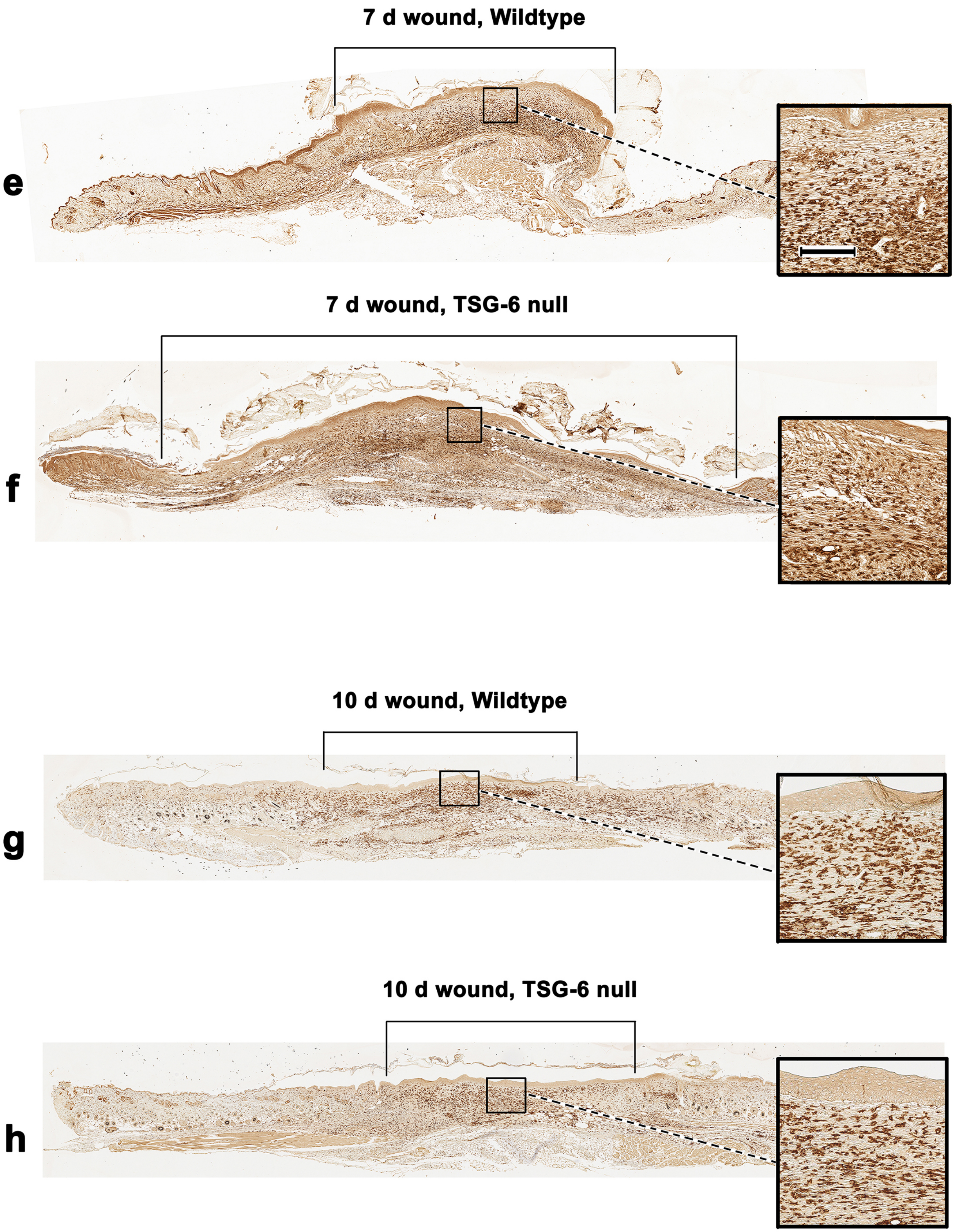

**Supplementary Figure S4**.

**Transition from granulation tissue to mature extracellular matrix in late skin wounds of wildtype and TSG-6 null mice.**

**Figure legend:**

Excisional skin wounds were harvested at 10 days post-injury, paraffin-fixed, and stained with Masson-Trichrome stain per the manufacturer's kit instructions. With this technique, the cytoplasm of cells stains red, nuclei stain dark red or black, and newly-formed collagen stains blue. Shown are three full-length wound beds from WT mice (panels a-c) or from TSG-6 null mice (panels d-f), with three enlargements of areas within the wound bed for each case. Note the greater cellularity and presence of blood vessels, and the lower amounts of collagen, in the TSG-6 null wounds as compared to the wildtype wounds. *Scale bar*, 100 μm.

**Supplementary Figure 4, panels a-c**

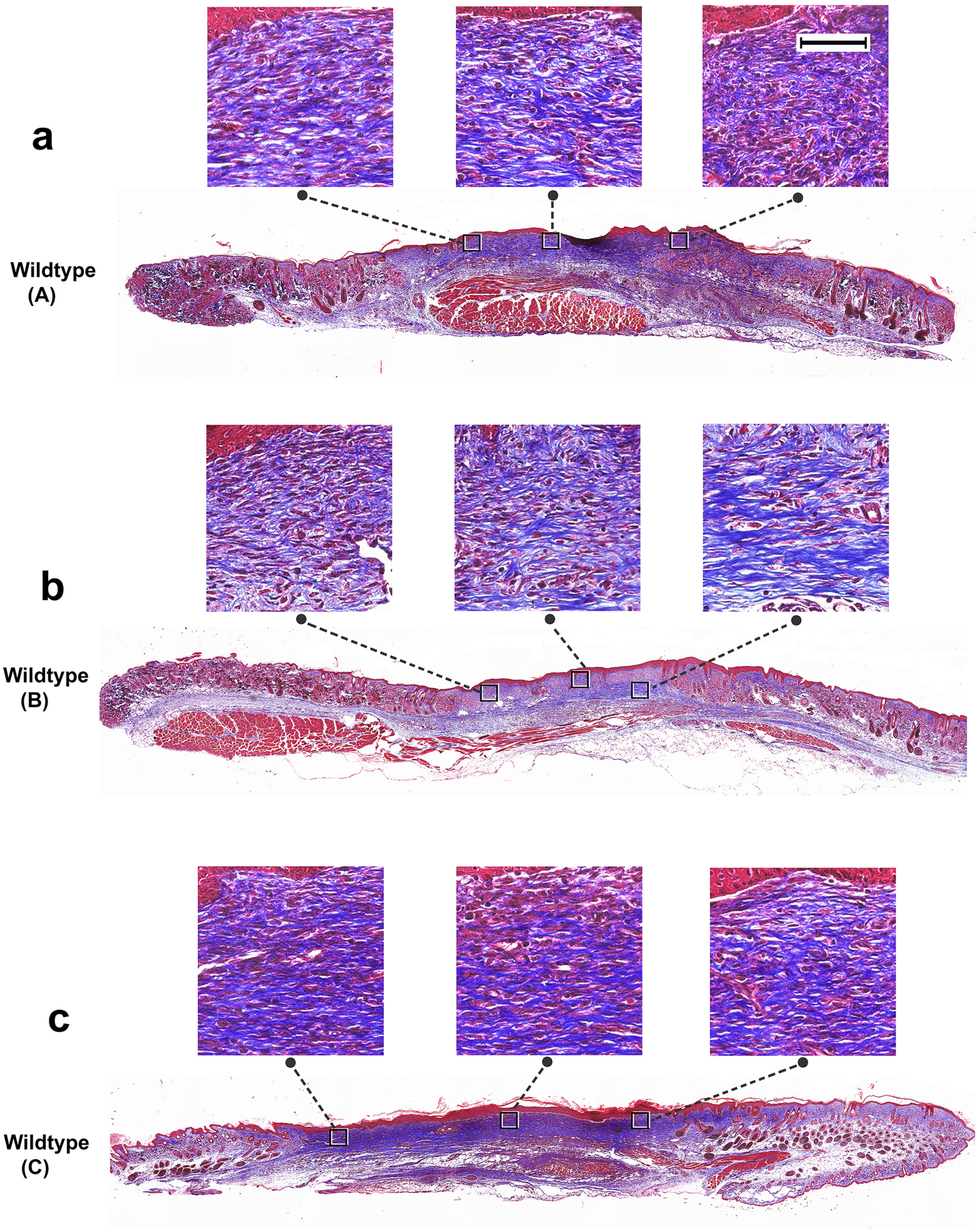

**Supplementary Figure 4, panels d-f**

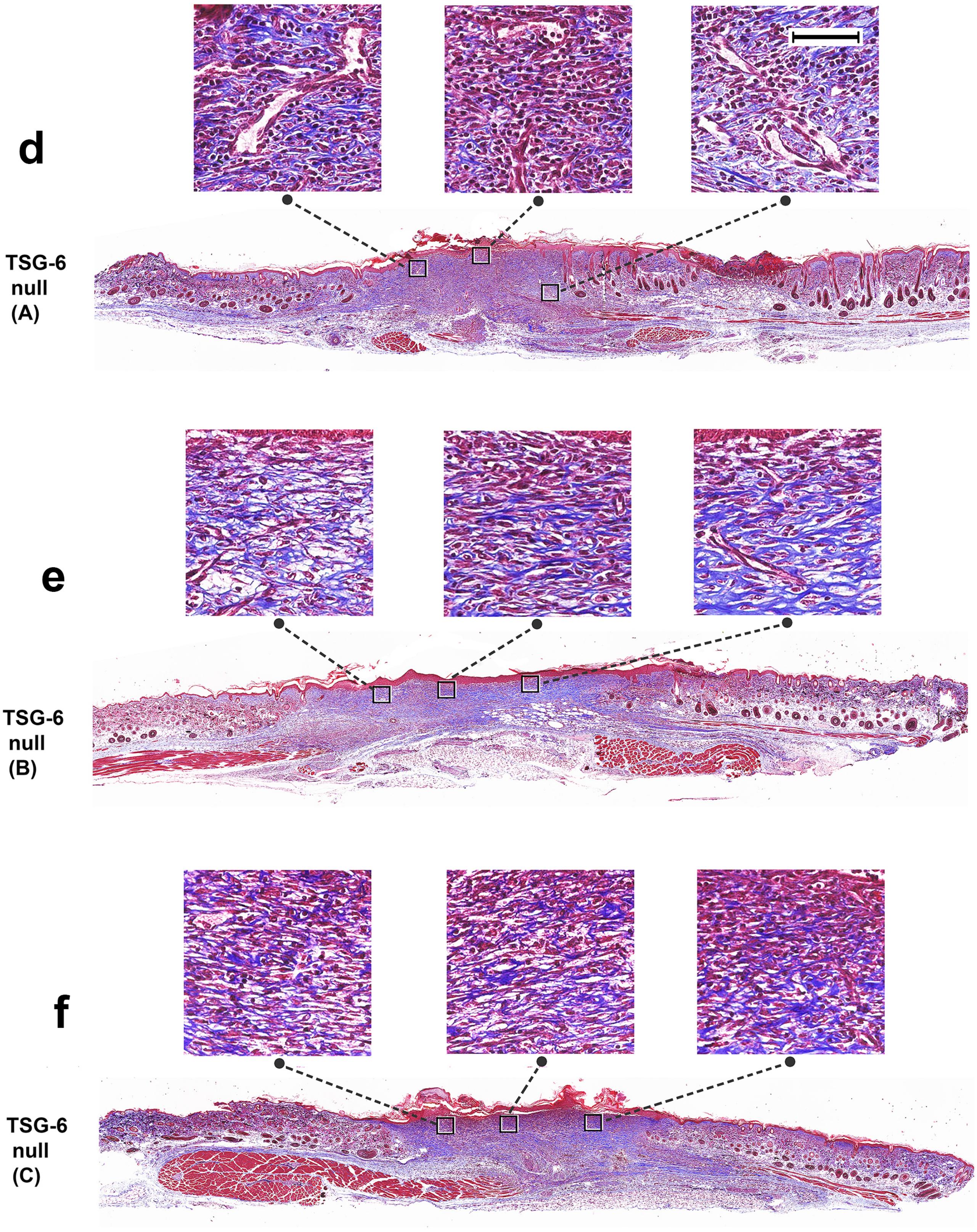

**Supplementary Figure S5.**

**Reintroduction of recombinant TSG-6 protein into TSG-6 null mouse wounds rescues the delayed closure phenotype.**

**Figure legend:**

Quantification of macroscopic wound closure between Days 0 to 7, for the experiment described in Figure 5 of the manuscript. Each excisional wound (5 mm diameter), in wildtype mice or in TSG-6 null mice, was injected with a total of 2 µg of rTSG-6 in 100 µL PBS, or with 100 µL of PBS vehicle alone. One injection was done on Day 0 immediately after wounding, and a second injection was administered on Day 4. Wounds were photographed and wound sizes expressed as percentage of initial area. Each bar represents the mean ± SEM; *(n)*, 8 wounds per condition. Statistical analysis was performed using Two Way ANOVA with Bonferroni correction; *P*-values are shown.

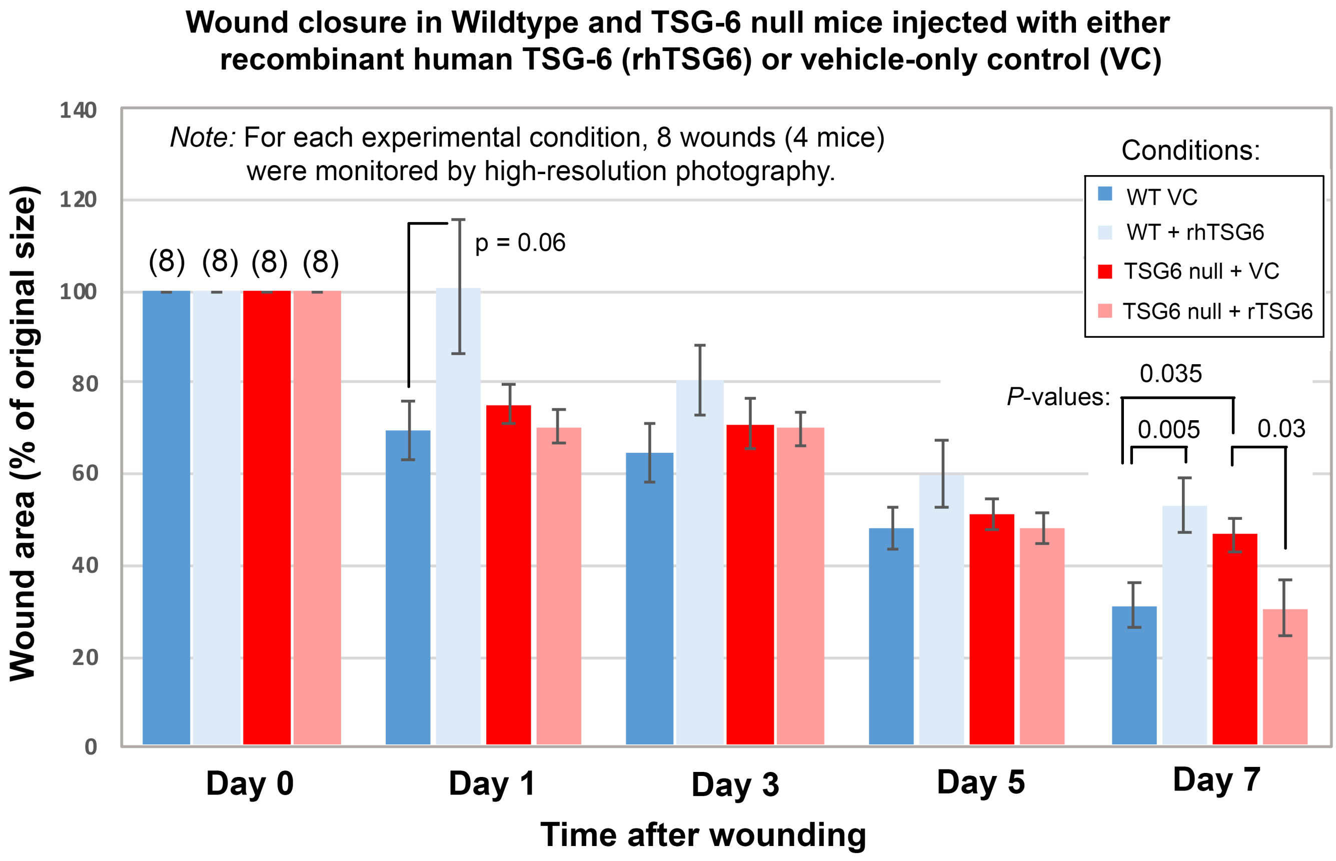

### Supplementary Figure S6.

**Analysis of cytokines in wildtype and TSG-6 null wounds.**

Cytokine protein assays were performed using a MILLIPLEX MAP magnetic bead panel for mouse cytokines/chemokines (Millipore/Sigma) on lysates from unwounded and wounded skin tissue. Excisional wounds were prepared using a 5 mm punch biopsy, and tissues were harvested at day 3, day 5, day 7, and day 10 after wounding. Unwounded skin (*No W*) was also collected. The sample solutions used in the assay had a protein concentration of 0.5 mg/ml. Data acquisition and analyses were done on the Luminex analyzer (MAGPIX), a CCD-based instrument. Multiple conjugated beads (internally color-coded microspheres with fluorescent dyes and coated with specific capture antibody) were added to each sample. The streptavidin-PE conjugate was used as a reporter molecule. The following graphs show concentrations of (**a**), IL-6; (**b**), KC; (**c**), MCP-1; (**d**), VEGF; and (**e**), IL-10. Mean ± SEM; (n) = number of mice; *p*-values from Kruskal-Wallis nonparametric tests.

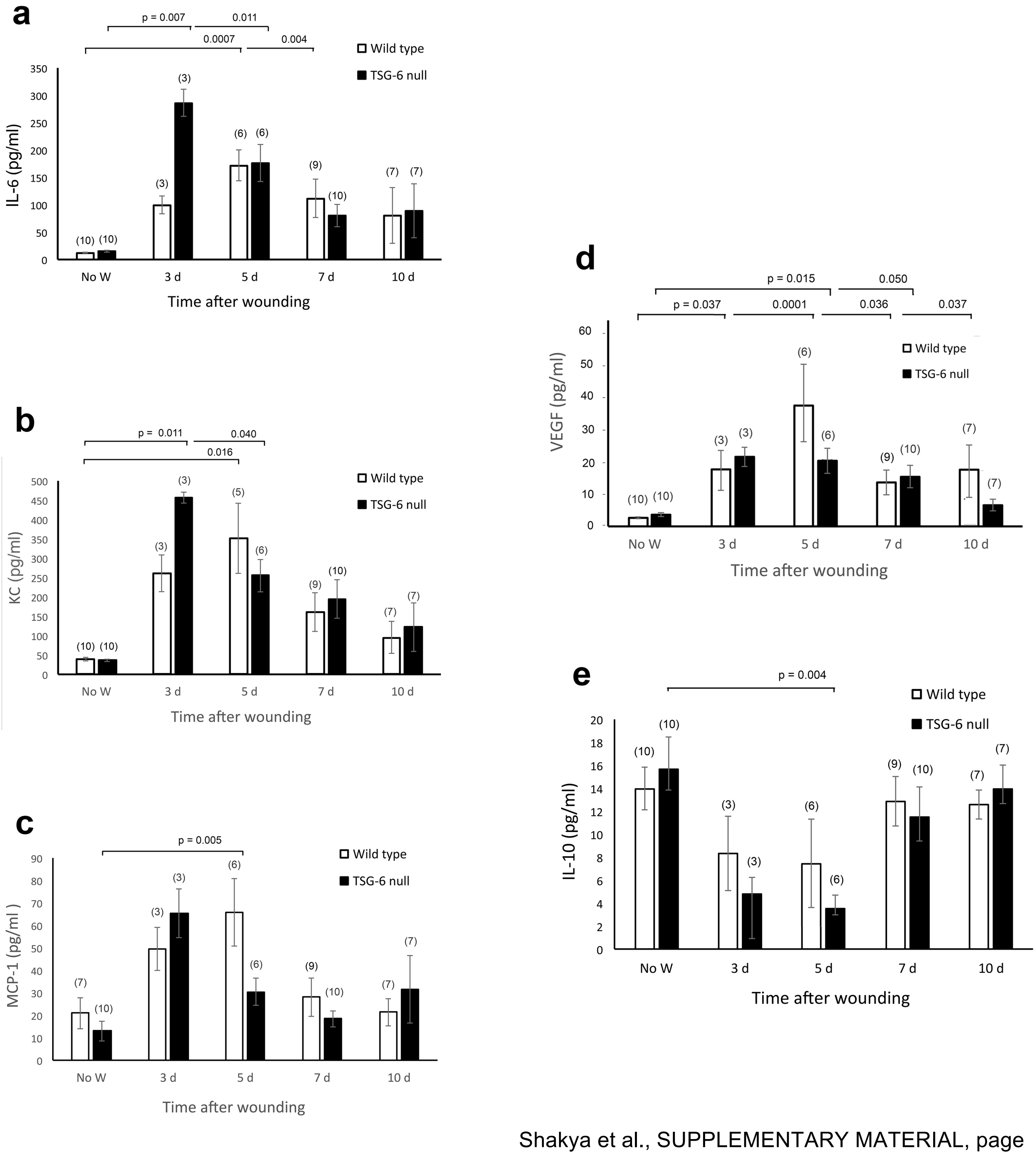
